# Weeds are not always evil: crop-weed relationships are context-dependent and cannot fully explain the positive effects of intercropping on yield

**DOI:** 10.1101/2020.04.02.021402

**Authors:** Laura Stefan, Nadine Engbersen, Christian Schöb

## Abstract

1. Implementing sustainable weed control strategies is a major challenge in agriculture. Intercropping offers a potential solution to control weed pressure by reducing the niche space available for weeds; however, available research on the relationship between crop diversity and weed pressure, and its consequences on crop yield is not fully conclusive yet.
2. In this study, we performed an extensive intercropping experiment using eight crop species and 40 different species mixtures to examine how crop diversity affects weed communities and how the subsequent changes in weeds influence crop yield. Field experiments were carried out in Switzerland and in Spain, which differ drastically in terms of climate and soil, and included monocultures, 2- and 4-species mixtures, and a control treatment without crops. Weed communities were assessed in terms of biomass, species richness and evenness, and community composition.
3. Results indicate that intercropping *per se* does not reduce weed performance or diversity, but crop species composition does. In particular, the presence of cereals in crop mixtures significantly reduced weed biomass and diversity.
4. Despite the lack of crop diversity effect on weeds, crop yield increased with crop species richness, particularly in Switzerland. Moreover, in Switzerland, where soil resources were abundant, increasing crop yield correlated with increased weed suppression and reduced weed diversity. In Spain, where water and nutrients were limited, crop yield was not related to weed biomass or diversity.
5. *Synthesis and applications*: We demonstrate that in our study, increased crop yield in mixtures was not due to increased weed suppression in diverse crop communities, but must be the result of other ecological processes. We also show that crop-weed relationships vary across environmental conditions, which emphasizes the need for a better understanding of weed communities’ assembly mechanisms, as well as locally adapted weed management strategies.

## 1. Introduction

Weeds are a major issue in agricultural systems, with the highest potential of crop yield loss compared to other pests (Oerke & Dehne, 2004). This is especially a problem in organic production, where the lack of herbicides forces farmers to adopt different weed control strategies (Bastiaans, Paolini, & Baumann, 2008). One option is to implement an ecological management of weeds (Mortensen, Bastiaans, & Sattin, 2000; Swanton & Weise, 1991). This strategy relies on the fact that crops and weeds compete for the same resources, such as water, nutrient, light, or space. If the pool of available resources cannot provide sufficient resources for both crops and weeds, competition between crops and weeds will arise, and any resource acquired by weeds will lead to a decrease in crop yield (Liebman, 1988). Therefore, by giving crops a head start and optimizing the use of resources by the crops, it is theoretically possible to reduce the available niche space for weeds, and consequently reduce weed pressure on crops.

Intercropping is an integrated crop management strategy that can be used to reduce weed pressure (Vandermeer, 1989). This is achieved through resource partitioning among co-occurring crop species with different resource acquisition strategies, allowing crops to better use the available resources, thereby decreasing the amount of resources available for weeds (Bybee-Finley, Mirsky, & Ryan, 2017; Pakeman et al., 2019). Resource partitioning is more likely to occur when functionally different crops are combined (Brun et al., 2019; Petchey & Gaston, 2002; Tilman et al., 1997). For instance, the combination of a cereal and a legume species, which differ in nitrogen acquisition, has been extensively studied and it has been shown to lead to higher yield and other beneficial services (Bedoussac et al., 2015; Connolly et al., 2017), such as decreased weed biomass in certain conditions (Bybee-Finley, Mirsky, & Ryan, 2017; Hauggaard-Nielsen, Ambus, & Jensen, 2001). Yet very few studies have examined the relationship between crop diversity and weed suppression more generally, i.e. using a wide variety of crop mixtures, combining more than two functional groups, or using combinations also differing from the cereal-legume combination (Baumann, Kropff, & Bastiaans, 2000; Szumigalski & Van Acker, 2005).

Traditionally, most weed research has focused on weed biomass, given the direct link between weed biomass and crop yield. Yet it remains unclear whether and how weed diversity and community composition are affected by crop species richness and functional composition, and how crop yield responds to such changes in weed diversity (Navas, 2012; Storkey & Neve, 2018). Previous research showed evidence for positive, negative and neutral effects of crop diversity on weed diversity (Palmer & Maurer, 1997; Schöb et al., 2017). In addition, how weed community composition changes with crop species richness remains unclear, in particular whether weed species colonizing diverse mixtures are a subset of the weeds in the species-poor crop communities, or if different weed species colonize the species-rich and species-poor crop communities respectively. Different weeds might indeed respond differently to the presence of a specific crop (Mohler & Liebman, 1987), and having a better understanding of which weed species grow in which crop mixtures could help pave the way to more targeted weed control strategies.

Efficient, sustainable weed control strategies also need to be adapted to local environmental conditions (Brooker et al., 2015); therefore, we need to understand how crop-weed relationships vary across different environmental conditions. It is for instance unclear how fertilization or climate influence the potential ability of intercropping to suppress weeds (Gomez & Gurevitch, 1998). Intercropping benefits are known to be usually higher in organic, low-productivity systems with little or no synthetic fertilizers (Bilalis et al., 2010; Brainard, Bellinder, & Kumar, 2011). Indeed, if resources are limited and fully exploited by crops, there should be little niche space available for weeds to grow. Therefore, we expect more weed suppression through intercropping in resource-poor, harsh environments.

Our research questions are the following: 1) How does crop diversity affect weed communities, in terms of biomass, diversity, and community composition? 2) Does crop productivity respond to the changes of the weed community? 3) How does the crop–weed interaction vary in different abiotic conditions? To answer these questions, we performed an intercropping experiment with eight different crop species belonging to four functionally different phylogenetic groups. We planted eight different monocultures, 24 different 2-species mixtures and 16 different 4-species mixtures in different climatic and fertilizing conditions, and examined the response of weed communities. We hypothesized that diverse intercrops will be better at suppressing weeds than monocrops, i.e. they would have fewer weed species and less biomass. We also expected that crop productivity and weed suppression (in terms of biomass and diversity) would be positively correlated, i.e. communities showing efficient weed suppression would have higher yield. Finally, we expected stronger weed suppression by crops under harsh abiotic conditions. In conclusion, our experiment comprehensively investigated crop–weed diversity–functioning relationships in varying abiotic conditions.

## 2. Material and methods

### 2.1 Study sites

The experiments took place in outdoor experimental gardens in Zurich, Switzerland, and in Torrejón el Rubio, Cáceres, Spain. In Zurich, the garden was located at the Irchel campus of the University of Zurich (47.3961 N, 8.5510 E, 508 m a.s.l.). In Torrejón el Rubio, the garden was situated at the Aprisco de Las Corchuelas research station (39.8133 N, 6.0003 W, 350 m a.s.l.). Zurich is characterized by a temperate climate, while the Spanish site is located in a dry, Mediterranean climate. The main climatic differences during the growing season are precipitation (572 mm in Zurich between April and August vs 218 mm in Cáceres between February and June) and daily average hours of sunshine (5.8 h in Zurich vs 8.4 h in Cáceres). Temperatures during the growing season do not vary substantially between the two sites: averages of daily mean, minimum and maximum temperatures are 14.0 °C, 9.3 °C and 18.6 °C in Zurich vs 14.6 °C, 9.6 °C and 19.6 °C in Cáceres. All climatic data are from the Deutsche Wetterdienst (www.dwd.de) and are average values over the years 1961 to 1990.

The experimental gardens were irrigated during the growing season with the aim of maintaining the above-mentioned differences in precipitation between the two sites while assuring survival of the crops during drought periods. In Spain, the irrigation was therefore configured for a dry threshold of soil moisture of 17% of field capacity, with a target of 25% of field capacity. In Switzerland, the dry threshold was set at 50% of field capacity, with a target of 90% of field capacity. Whenever dry thresholds were reached (measured through PlantCare soil moisture sensors (PlantCare Ltd., Switzerland)), irrigation started and added water up to the target value.

Each experimental garden consisted of 392 square boxes of 0.25 m^2^ with a depth of 40 cm. Beneath 40 cm the boxes were open, allowing unlimited root growth. The boxes were embedded into larger beds: in Switzerland, there were 14 beds of 14×1 m^2^, each bed containing 28 plots. In Spain, beds were 20×1 m^2^ and contained 40 plots. Inside a bed, plots were separated from each other by metal frames. Each box was filled until 30 cm with standard, not enriched, agricultural soil coming from the local region. Soil structure and composition therefore varied between sites and reflect the environmental histories of each site. In Spain, soil was composed of 78% sand, 20% silt, and 2% clay, and contained 0.05% nitrogen, 0.5% carbon, 253 mg total P/kg. In Switzerland, the soil consisted of 45% sand, 45% silt, and 10% clay, and contained 0.19% nitrogen, 0.42% carbon, and 332 mg total P/kg. Swiss soils had a mean pH of 7.25, while in Spanish soils the pH was 6.30.

We fertilized half of the beds with nitrogen, phosphorus and potassium at the concentration of 120 kg/ha N, 205 kg/ha P, and 120 kg/ha K. Fertilizers were applied three times, namely once just before sowing (50 kg/ha N), once when wheat was at the tillering stage (50 kg/ha N), and once when wheat was flowering (20 kg/ha N). The other half of the beds served as unfertilized controls. We randomly allocated individual beds to a fertilization or non-fertilized control treatment.

### 2.2. Crop species and cultivars

At each site, experimental communities were constructed with eight annual crop species. We used crop species belonging to four different phylogenetic groups with varying functional characteristics: *Triticum aestivum* (wheat, C3 grass) and *Avena sativa* (oat, C3 grass), *Lens culinaris* (lentil, legume) and *Lupinus angustifolius* (lupin, legume), *Linum usitatissimum* (flax, herb [superrosids]) and *Camelina sativa* (false flax, herb [superrosids]), and *Chenopodium quinoa* (quinoa, herb [superasterids]) and *Coriandrum sativum* (coriander, herb [superasterids]). This resulted in species combinations with different phylogenetic distances: Cereals diverged from the three other groups 160 million years ago; superasterid herbs then diverged from superrosid herbs and legumes 117 million years ago. Finally, legumes diverged from superrosid herbs 106 million years ago (timetree.org). We chose phylogenetic distance as a criterion for functional similarity as it is often positively correlated with functional diversity and thus a good proxy to assess the impacts of species diversity on ecosystem functions (Mouquet *et al*., 2015). Furthermore, we chose crop cultivars that were locally adapted to each site (Table S1 in SI) and replicated the experimental setup with both Spanish and Swiss cultivars at each site.

### 2.3 Experimental crop communities

Experimental communities consisted of control plots with no crops, monocultures, 2- and 4-species mixtures. We planted every possible combination of 2-species mixtures with two species from different phylogenetic groups and every possible 4-species mixture with a species from each of the four different phylogenetic groups present. We replicated the experiment two times with the exact same species composition. Plots were randomized within each country, fertilizer and ecotype (i.e. cultivar origin) treatment. Each monoculture and mixture community consisted of one, two or four species planted in four rows. Two species mixtures were organized following a speciesA|speciesB|speciesA|speciesB pattern. The order of the species was chosen randomly. For 4-species mixtures, the order of the species was also randomized. Density of sowing differed among species groups and was based on current cultivation practices: 160 seeds/m^2^ for legumes, 240 seeds/m^2^ for superasterids, 400 seeds/m^2^ for cereals, and 592 seeds/m^2^ for superrosids. We sowed by hand in early February 2018 in Spain and early April 2018 in Switzerland.

### 2.4 Data collection

We assessed weed biomass and species richness in each plot at the flowering stage of the crops (May 2018 in Spain, 85 days after sowing; June 2018 in Switzerland, 70 days after sowing). Weeds were cut at the soil surface, divided by species, dried in the oven at 80°C until constant weight, and then weighed.

Grain yield of each crop species was determined in each plot when fruits were ripe. This corresponded to mid-June/July in Spain and July/August in Switzerland. As time of maturity slightly varied among the different crops, we conducted harvest species by species. In Spain, we first harvested *Lupinus* 135 days after sowing, followed by *Avena, Triticum, Coriandrum* and *Lens* 145 days after sowing, *Camelina* 150 days after sowing, *Linum* 160 days after sowing, and *Chenopodium* 183 days after sowing. In Switzerland, we first harvested *Camelina* 100 days after sowing, followed by *Avena* and *Triticum* 115 days after sowing, *Coriandrum, Lens* and *Lupinus* 130 days after sowing, *Linum* 140 days after sowing, and *Chenopodium* 150 days after sowing. We clipped plants right above the soil surface and separated seeds from the vegetative parts. Seeds were sun-dried for five days and weighed.

### 2.5 Data analyses

For data analyses, we eliminated plots with incomplete data. Due to birds foraging on wheat seeds, we discarded a substantial number of plots containing wheat in Spain. Final counts were 294 plots in Spain, 328 in Switzerland, for a total of 622 plots.

For each plot, a Weed Suppression Index was calculated using 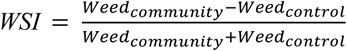, with *Weed*_*community*_ being the total weed biomass collected in a plot with crops, and *Weed*_*control*_ being the average of total weed biomass collected in control plots without crops for each combination of country and fertilizer treatments. A negative WSI means that there is less weed biomass in the community plots compared to the controls. A WSI close to -1 indicates that the weed suppression effect is high, i.e. that there are almost no weeds present in the plot.

We calculated Shannon’s diversity (H) and evenness (J) indexes for the weed community of each plot.

We used linear mixed models to analyze the effects of the experimental treatments on WSI, weed species number, H, and J, respectively. Fixed factors included country, fertilizing condition, ecotype, crop species number, presence of cereal, legume, superrosid or superasterid herb, as well as the interactions between them (except interactions between crop species number and presence of functional groups). Species composition and bed were set as random factors.

Responses of the weed community composition to the experimental treatments was analyzed using permutational multivariate analyses of variance and distance-based redundancy analyses with Bray-Curtis distance (Legendre & Andersson, 1999). Due to a high variability in weed species between Spain and Switzerland, we analyzed weed communities in the two countries separately. To that end, in each country, we removed rare species (i.e. weeds present in only one or two plots), and log-transformed weed biomass. We first performed permanova tests with Bray-Curtis distance, using the function a*donis* from the *vegan* package in R (Anderson, 2001). Experimental factors tested included environmental factors (country, fertilizer), ecotype, crop diversity treatments (crop species number, presence of the different functional groups), and their interactions. We then performed constrained ordinations with Bray-Curtis distance using the functions *capscale* from the vegan package in R (Legendre & Andersson, 1999).

Total crop yield per plot was calculated, transformed using Tukey’s ladder of power (transformation to the power of 0.2) (Tukey, 1957) and analyzed using a linear mixed model, with species combination and bed as random factors, and with fixed factors including the ones described before, as well as WSI, weed species number, and their interactions with country and fertilizer. Pairwise multiple comparisons using post-hoc Tukey tests were performed to test for the effects of species number within each country.

## 3. Results

### 3.1 Weed Suppression Index

Weed Suppression Index (WSI) varied according to country, fertilizer, cereal presence, ecotype, country×fertilizer, country×cereal, country×legume, and country×fertilizer×cereal (Table S2 of SI). Country had the strongest effect on WSI: WSI was 52% lower in Switzerland than in Spain (Fig. 1a). Fertilizer application reduced WSI by 24%, while the presence of a cereal led to a reduction of 39%. WSI was lower in plots containing Swiss ecotypes compared to Spanish ecotypes (−12%) (Fig. 1b) Cereal presence decreased WSI more in Spain (− 84%) compared to Switzerland (−12%) (Fig. 1c). The effect of fertilizer application was also stronger in Spain (− 50%) than in Switzerland (−13%). Legume presence slightly decreased WSI in Spain (−2%), while it increased WSI in Switzerland (+6%). Crop species number had no significant effect on WSI.

**Fig 1.**
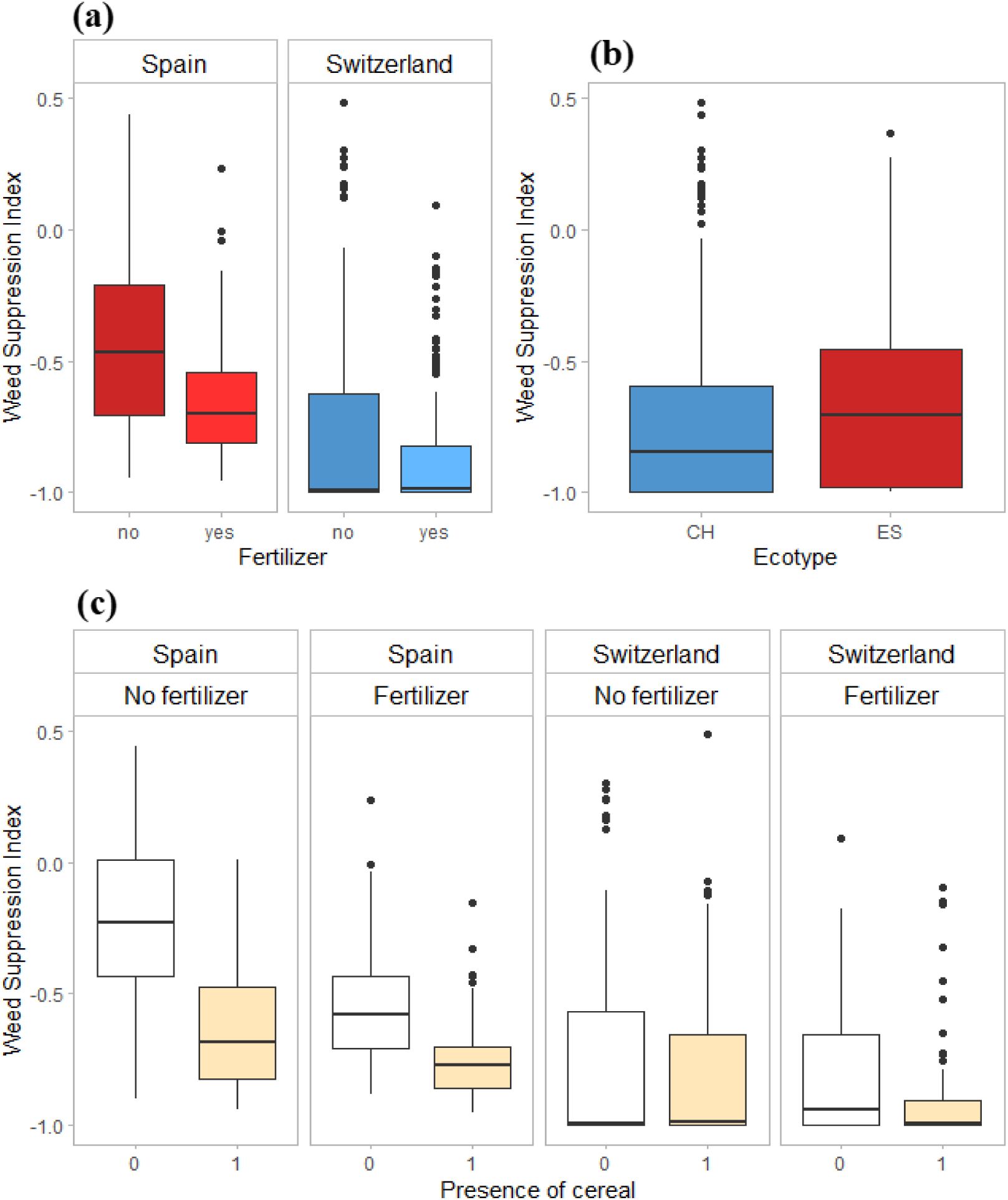
Effects of country and fertilization (a), ecotype (b), and presence of a cereal (c) on Weed Suppression Index. (Abbreviations: CH = Swiss ecotype, ES = Spanish ecotype).

### 3.2 Weed diversity (species number, Shannon diversity, Shannon evenness)

Weed species richness was significantly influenced by country, fertilizer, ecotype, country×cereal, country×legume, and country×superasterid herb (Table 1, Fig S1 of SI). Weed species richness was lower in Switzerland compared to Spain (−57%), in unfertilized plots compared to fertilized plots (−16%), and in Swiss ecotype plots compared to Spanish ecotypes (−10%). Cereal presence decreased weed species richness in Spain by 6%, but had no effect in Switzerland. Legume presence increased weed species richness in Switzerland by 20% but had no effect in Spain. Finally, the presence of a superasterid herb increased weed species richness in Spain (+1%), but decreased it in Switzerland (−3%).

**Table 1.** Results of mixed effects ANOVA testing environmental factors, crop diversity, ecotype and functional group ence on weed species richness, Shannon’s diversity index (H), and Shannon’s evenness index (J). p-values in bold are significant at α = 0.05; * (P < 0.05), ** (P < 0.01), *** (P < 0.001).

|  | *Weed species richness*<br>*F value* | *Weed species richness*<br>*p-value* | *H*<br>*F value* | *H*<br>*p-value* | *J*<br>*F value* | *J*<br>*p-value* |
| --- | --- | --- | --- | --- | --- | --- |
| *Country* | 356.5336 | **< 2.2e-16 \*\*\*** | 6.89E+02 | **< 2e-16 \*\*\*** | 243.4939 | **8.673e-13 \*\*\*** |
| *Fertilizer* | 12.5977 | **0.001578 \*\*** | 4.96E+00 | **0.03523 \*** | 1.0365 | 0.32066 |
| *Crop species number* | 0.5227 | 0.59692 | 0.8429 | 0.43839 | 1.7064 | 0.18252 |
| *Country x fertilizer* | 0.0375 | 0.848063 | 1.24E+00 | 0.27597 | 3.5899 | 0.07238 |
| *Country x crop sp.number* | 0.1108 | 0.895109 | 0.1635 | 0.84919 | 1.5433 | 0.21463 |
| *Fertilizer x crop sp.number* | 0.405 | 0.667199 | 0.1299 | 0.87822 | 0.0819 | 0.92137 |
| *Country x fertilizer x crop sp.number* | 0.9321 | 0.394336 | 2.3001 | 0.10121 | 1.3821 | 0.25198 |
| *Cereal* | 2.2359 | 0.142466 | 3.3143 | 0.07621 | 0.3422 | 0.55883 |
| *Legume* | 0.3871 | 0.537412 | 0.3126 | 0.57948 | 0.6997 | 0.40326 |
| *Superasterid herb* | 0.0716 | 0.790408 | 0.0912 | 0.76431 | 0.522 | 0.47032 |
| *Ecotype* | 10.3945 | **0.001338 \*\*** | 4.2982 | **0.03861 \*** | 0.2019 | 0.65334 |
| *Country x cereal* | 6.3491 | **0.012022 \*** | 6.13E+00 | **0.01357 \*** | 1.0331 | 0.3099 |
| *Fertilizer x cereal* | 3.8083 | 0.051509 | 2.5657 | 0.10978 | 1.454 | 0.22843 |
| *Country x legume* | 20.1559 | **8.65E-06 \*\*\*** | 0.4853 | 0.48633 | 0.2809 | 0.59636 |
| *Country x ecotype* | 1.2864 | 0.257192 | 0.0171 | 0.89602 | 0.0084 | 0.92708 |
| *Fertilizer x legume* | 2.4956 | 0.114737 | 0.8462 | 0.35802 | 0.1069 | 0.74387 |
| *Country x superasterid* | 7.8013 | **0.005399 \*\*** | 3.9231 | **0.04811 \*** | 1.2838 | 0.25769 |
| *Fertilizer x superasterid* | 0.3367 | 0.561991 | 0.8689 | 0.35166 | 0.0003 | 0.98712 |

Shannons’s diversity index (H) was likewise affected by country, fertilizer, ecotype, country×cereal and country×superasterid herb (Table 1, Fig S1 of SI). Similarly to weed species richness, H was lower in Switzerland (−75%), in unfertilized plots (−12%), and with Swiss ecotypes (−11%). Cereal presence had a positive effect on H in Switzerland (+9%), while in Spain the effect of cereal was slightly negative (−5%). Presence of a superasterid herb increased H in Switzerland (+22%) but not in Spain.

Shannon’s evenness index (J) was only affected by country (Table 1, Fig S1 of SI): J was lower in Switzerland (−58%) compared to Spain.

Crop species number had no significant effects on any of the weed diversity metrics.

### 3.3 Weed community composition

In Spain, weed communities were dominated by *Poa annua* and *Spergula arvensis* in terms of biomass, and by *Poa annua, Arabidopsis thaliana* and *Spergula arvensis* in terms of frequency (see Table S3 in SI for a complete weed species list). Evenness was generally high, with an average of 0.64, meaning an even distribution of biomass across weed species. Weed community composition was significantly affected by fertilizer (p = 0.001, Fig. 2b), crop species number (p = 0.027, Fig 2d), cereal presence (p = 0.001, Fig 2f), legume presence (p = 0.029), superasterid presence (p = 0.042), ecotype (p = 0.001), fertilizer×cereal (p = 0.001) and crop species number×cereal (p = 0.005, Fig S2b in SI) (Table S3 in SI). Ordination plots show that diversity treatments are nested into each other in terms of weed community composition, i.e. the composition of weed communities in the 4-species mixtures is a subset of the weed composition in the 2-species mixtures, which is itself embedded in the monocultures (Fig 2b).

**Fig 2.**
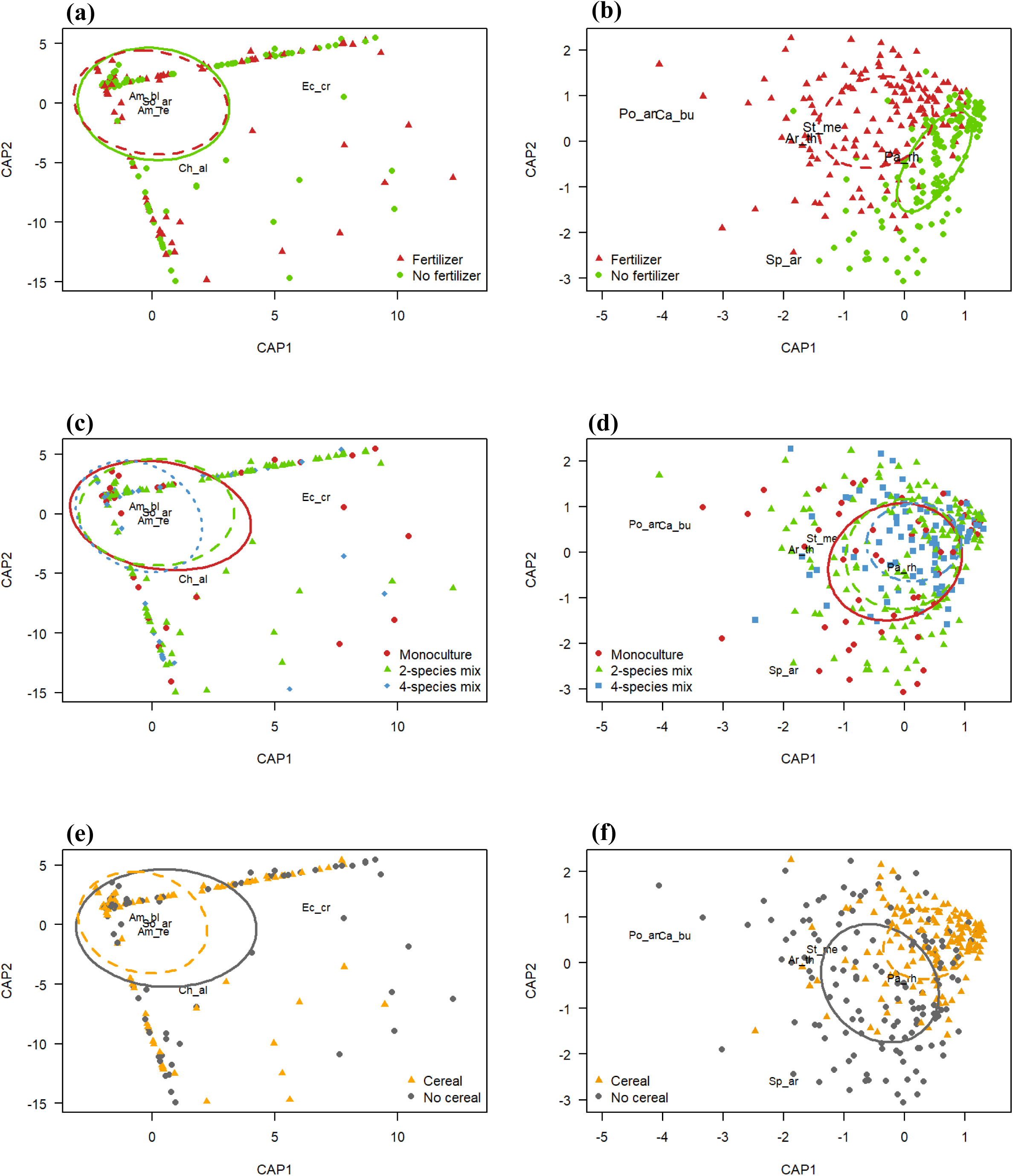
Constrained ordination plots showing changes in weed communities in Spain (right panel) and Switzerland (left panel) in ponse to fertilization (a-b), crop species number (c-d), and presence of cereal (e-f). The most responsive weed species are also resented (abbreviations: Po_an: *Poa annua;* St_me: *Stellaria media;* Pa_rh: *Papaver rhoeas;* Ar_th: *Arabidopsis thaliana;* _ar: *Spergula arvensis;* Cap_bu: *Capsella bursa-pastoris;* Ec_cr: *Echinocloa crus-galli;* Ch_al: *Chenopodium album;* Am_re: *aranthus retroflexus;* Am_bl: *Amaranthus blithum;* So_ar: *Sonchus arvensis.*).

In Switzerland, weed communities were dominated by *Chenopodium album* and *Echinocloa crus-galli* in terms of biomass, and by *Sonchus arvensis* and *Amaranthus retroflexus* in terms of frequency (see Table S3 in SI for a complete weed species list). Evenness was low, with an average of 0.27, which means that biomass of weed species was unequally distributed across the weed species. Results of the permutational analysis showed that weed communities were affected by fertilizer (p = 0.001, Fig 2a) and ecotype (p = 0.009) (Table S3 in SI). Crop species number did not significantly affect weed community composition; however, graphical representation of the ordination shows that the 4-species mixtures are nested in the two-species mixtures and monocultures (Fig. 2c).

### 3.4 Crop yield

Total crop yield per plot was significantly affected by weed suppression index, weed species richness, fertilizer, crop species richness, ecotype, presence of superasterid herb, country×WSI, country×weed species number, fertilizer×weed species number, country×crop species number, country×cereal, and country×legume (Table S4 in SI). Overall, WSI and weed species richness had a negative effect on total crop yield (slope for WSI = -100.88, slope for weed species richness = -18). However, this effect was significantly stronger in Switzerland (slope for WSI = -113.5, slope for weed species richness = -26.2) compared to Spain (slope for WSI = -16, slope for weed species richness = -1.8) (Fig. 3). Fertilizer increased crop yield by only 2%. Crop species richness significantly increased crop yield (Fig. 4a): 2-species mixtures had 29% more crop yield than monocultures, and 4-species mixtures had on average 53% increase in yield compared to monocultures. Pairwise comparisons however showed that this effect was only significant in Switzerland, where 2-species mixtures and 4-species mixtures had 35% and 78% more crop yield than monocultures, respectively. Yield was higher with Swiss ecotypes (+71%) compared to Spanish ecotypes. Cereal presence decreased crop yield by 6% in Spain, while it increased yield by 54% in Switzerland (Fig. 4b). Legume presence, on the contrary, increased crop yield in Spain (+68%) but decreased yield in Switzerland (−24%) (Fig 4c). Superasterid herb presence slightly decreased crop yield (− 2.5%).

**Fig. 3.**
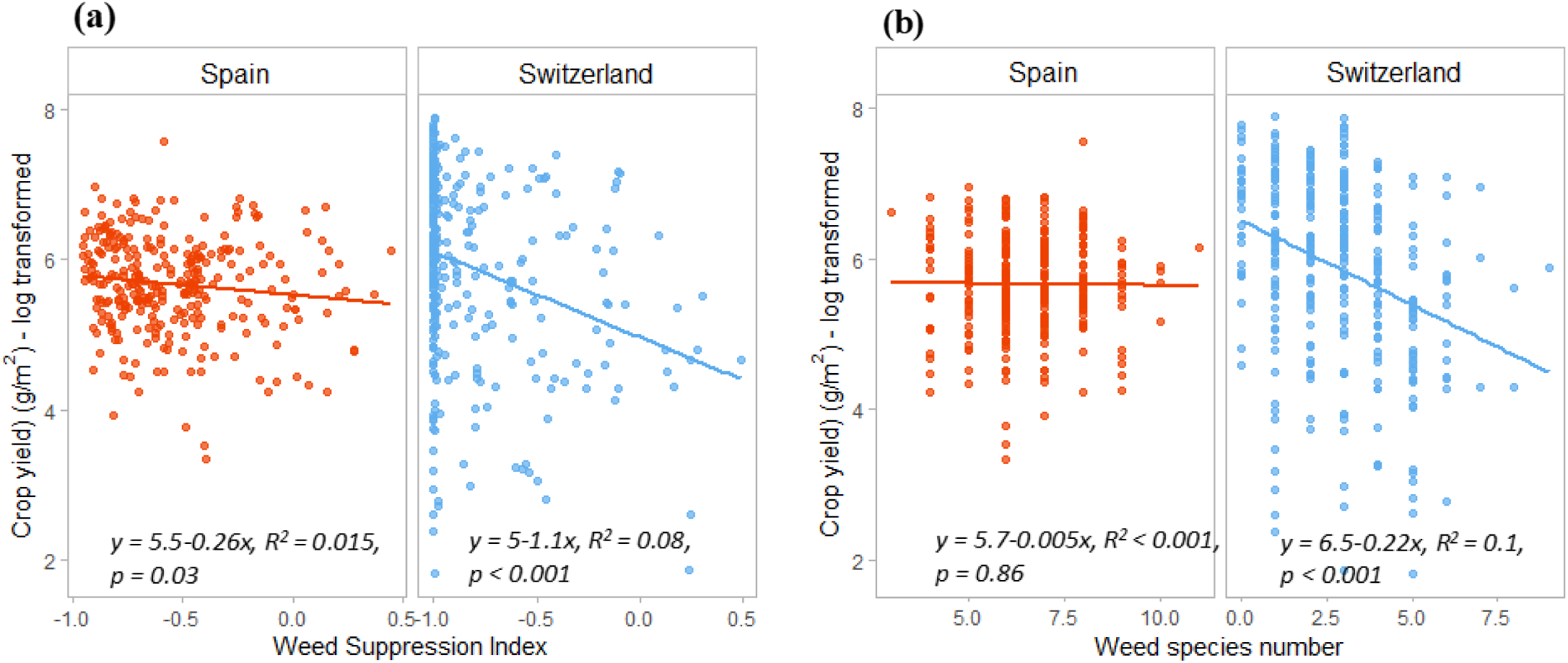
Relationship between crop yield and Weed Suppression Index (a) and weed species number (b) in Spain and Switzerland.

**Fig. 4.**
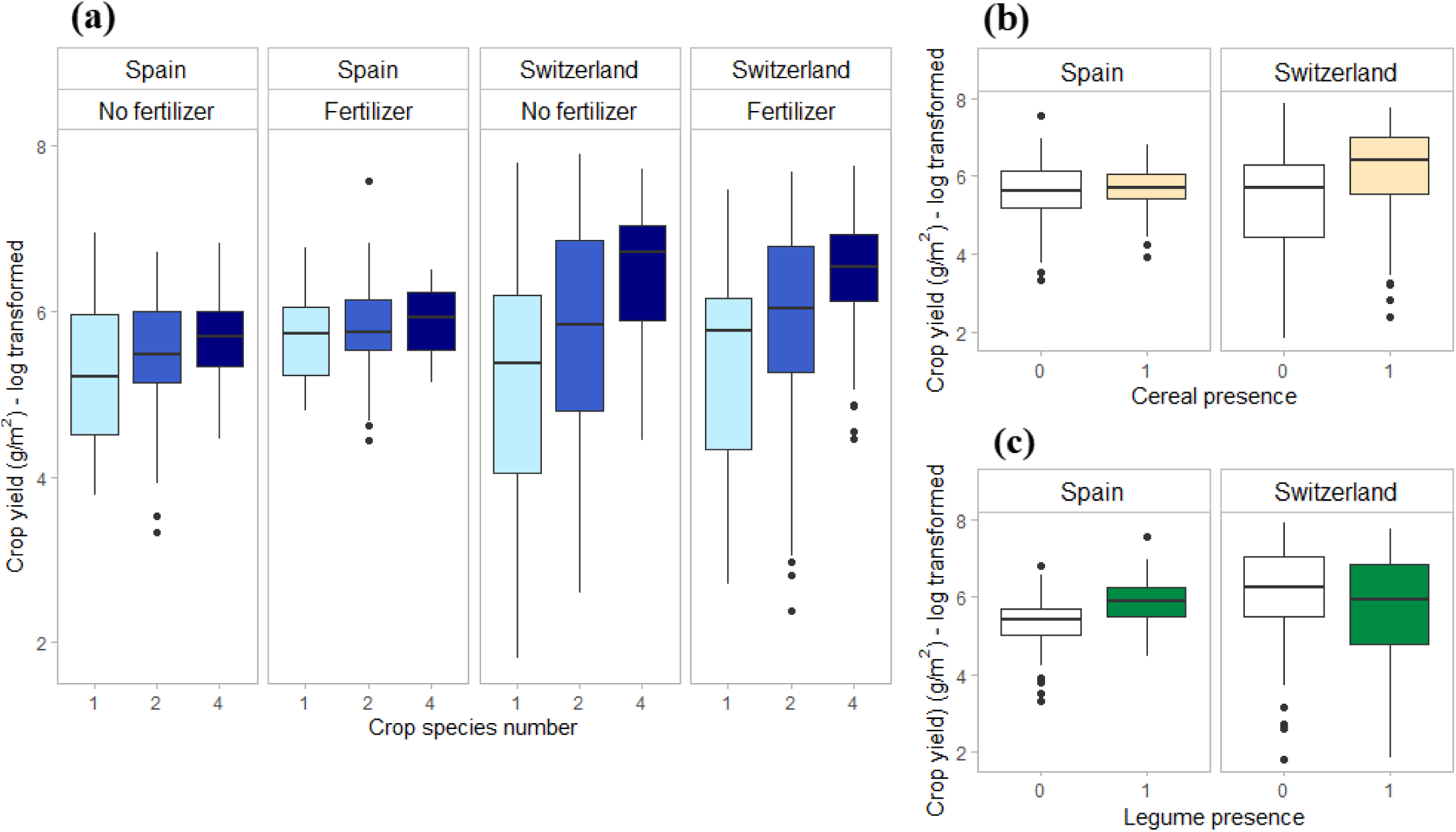
Effects of crop species number (a), cereal presence (b), and legume presence (c) on crop yield, in Spain and in Switzerland.

## 4. Discussion

Our study shows no significant effect of crop species number on weed biomass nor weed species richness, but a change in the weed community composition, with the weed community in diverse mixtures being mainly a subset of the weed communities in less diverse crop communities. Despite the lack of a crop diversity effect on weeds, crop mixtures showed increased yield compared to monocultures, in particular in Switzerland. Furthermore, stronger weed suppression and reduced weed diversity were correlated with increased crop yield in Switzerland. This demonstrates that in our study, increased crop yield in mixtures was not driven by increased weed suppression of diverse crop communities but must be attributed to other ecological processes.

### Crop species number effects on weeds

We hypothesized that increasing crop diversity would enhance weed suppression and decrease weed diversity due to a more complete use of the available niche space by the crops. This was not the case in our experiment: crop species richness had no significant effect on the Weed Suppression Index or on any measure of weed diversity. This suggests that crop mixtures do not necessarily suppress weeds more than monocultures. This result is consistent with some previous studies showing that intercropping *per se* does not affect weed biomass while crop identity and mixture composition does (Bybee-Finley, Mirsky, & Ryan, 2017). Indeed, we observed a strong effect of cereals on WSI, demonstrating that cereals are very performant at suppressing weeds (Fig 1b). This is coherent with many studies looking at cereal-legume intercrops, where the intercrop often reduces weed biomass compared to the sole legume but not the cereal monocrop (Corre-Hellou et al., 2011; Poggio, 2005; Szumigalski & Van Acker, 2005). We also observed a negative effect of cereal presence on Shannon’s diversity index, which suggests that cereals not only reduce weed performance, but also limit weed diversity. Furthermore, cereal presence had a strong effect on weed community composition in Spain (Fig 3c). Cereals are commonly considered to be strong competitors, due to their high growth rate at early life stages, their extensive rooting system, and their high uptake of nitrogen (Andrew, Storkey, & Sparkes, 2015; Bertholdsson, 2005; Ofori & Stern, 1987). Moreover, some species of cereals, including wheat, present allelopathic abilities that have been demonstrated to be of importance in increasing competitiveness against weeds (Jabran, Mahajan, Sardana, & Chauhan, 2015; Worthington & Reberg-Horton, 2013).

### Weed effects on crop yield

Our second goal was to investigate whether weed responses would correlate with changes in crop yield. Specifically, we expected that higher weed pressure would relate to a reduction in yield. Overall, when considering all abiotic conditions, we did find that crop productivity was positively correlated with weed suppression. Total crop yield was higher in plots with lower weed species richness and higher weed suppression ability. This result is consistent with many studies (e.g. Bybee-Finley, Mirsky & Ryan, 2017; Adeux *et al.*, 2019), which also find a negative relationship between crop productivity and weed biomass, and it suggests that crops and weeds compete for the same resources and hinder each other’s performances. The effects of our ecotype treatment on crops and weeds supports this idea: the Swiss ecotypes showed higher yield, but also lower weed diversity and biomass compared to the Spanish varieties (Fig S1 of SI, Table 2). Crop yield was also positively linked to crop species richness, showing a positive diversity–productivity relationship (Fig 4a). However, since weed performance was not affected by crop species number, the beneficial effects of crop species richness on yield does not seem to be due to a reduction in weed pressure. Rather, we suggest that other ecological processes must play a role in increasing crop productivity in diverse mixtures, such as nutrient partitioning, (Jensen, Carlsson, & Hauggaard-Nielsen, 2020; Von Felten et al., 2009), light partitioning (Jesch et al., 2018; Spehn et al., 2005), or changes in belowground microbial communities (Duchene, Vian, & Celette, 2017; Lange et al., 2015). We therefore think that higher crop productivity driven by these other factors enhances competition between crops and weeds and suppresses weeds, rather than the other way around.

### The environmental context-dependence of crop–weed interactions

The effects described above differed in strength and direction in different abiotic conditions. Indeed, there was a strong effect of country, fertilizer, and their interaction, on WSI, weed species richness, H, and weed community composition. First, we found that in Switzerland, crops were more performant at reducing weeds and weed communities were less diverse. Switzerland has a temperate climate, while Spain is semi-arid; this difference in climate, combined with the restricted irrigation and the lower soil nitrogen in Spain suggest that both crops and weeds experienced harsher abiotic conditions in Spain. Contrary to our third hypothesis, weed suppression by crops was not higher under such limiting environmental conditions. Furthermore, WSI was lower in fertilized conditions, which means that crops reduce weed biomass more efficiently in fertilized plots compared to unfertilized plots. This suggests more and stronger competition, and therefore increased weed suppression by crops, in mild environments compared to harsh abiotic conditions (Fig 5). This is consistent with the stress-gradient hypothesis, suggesting stronger and more frequent competition in non-stressful environments (Bertness & Callaway, 1994).

**Fig. 5.**
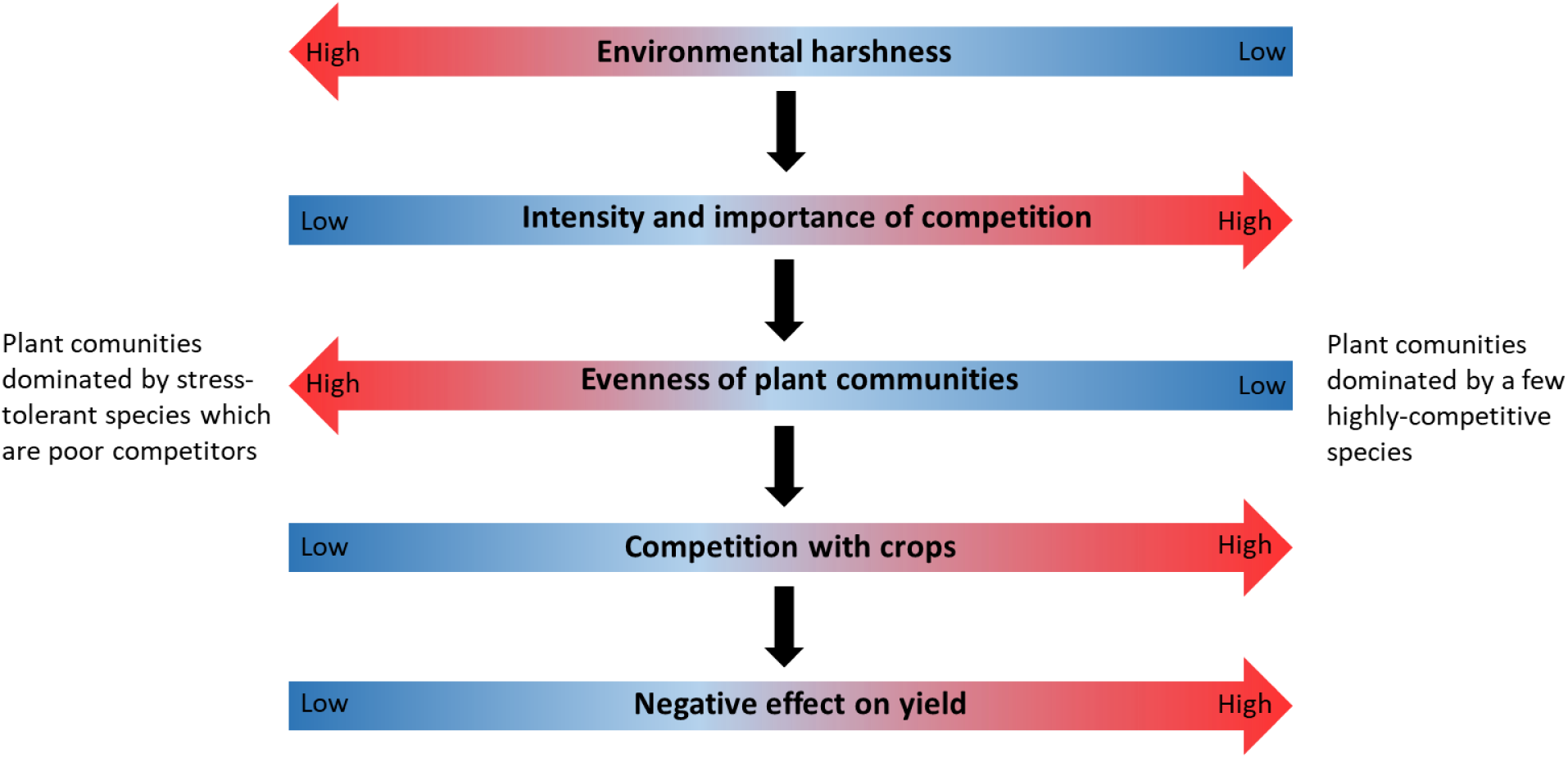
Conceptual figure depicting the context-dependence of crop-weed relationships.

Secondly, the effect of weeds on crop yield also varied among countries (Fig 3). In Spain, crop yield was not affected by WSI nor weed species richness (Fig 3a). Given that the Spanish communities had a significantly lower yield compared to Switzerland, our results thus suggest that in low-productive systems, crop productivity may not be affected by weeds. This hypothesis is reinforced by the effect of cereal presence in Spanish communities: despite the fact that cereal presence decreases weed biomass, crop yield does not subsequently increase. Therefore, crop productivity does not seem to depend on weed pressure in Spain. It does however in Switzerland, where yield is negatively affected by WSI and weed species richness (Fig 3b). In addition, contrary to Spain, the beneficial effect of cereal presence on weeds actually translates into an increase in yield. Hence, in Switzerland, more weeds correlate with lower crop yield. This supports our previous statement that in mild environments, crops and weeds strongly compete with each other, while in low-productive, resource-poor systems competitive interactions are less important, potentially because plant biomass and consequently resource depletion is lower (Goldberg & Novoplansky, 1997; Grime, 1973, 1979). This would also explain why in Spain weeds mainly respond to abiotic conditions and crop composition - as illustrated by the results of weed community composition - while in Switzerland they correlate with crop productivity independently of environmental conditions.

Changes in competition intensity and importance are known to influence plant community structure (Tilman, 1982); in particular, it has been reported that increasing competition intensity and importance can be associated with a decline in evenness (Lamb & Cahill, 2008; Lamb, Kembel, & Cahill, 2009). In our study, weed communities in Spain indeed had a higher evenness and higher species richness than in Switzerland, where two very competitive species - *Chenopodium album* and *Echinochloa crus-galli* – were completely dominant. These two weed species thrive on nitrogen-rich soils, particularly *Echinochloa*, which can remove available soil nitrogen by up to 80% and thereby significantly reduce crop yield (Randall, 2012; Heuzé, 2017). Weed evenness can consecutively influence crop yield: increasing weed community evenness decreases weed biomass and increases crop productivity (Adeux *et al*. 2019). It is therefore suggested that more diverse and even weed communities could limit the negative effects of dominant, highly-adapted, competitive weed species on crop yield (Cierjacks, Pommeranz, Schulz, & Almeida-Cortez, 2016; Storkey & Neve, 2018). Henceforth, we hypothesize that in harsher, unproductive environments, competitive interactions are less important, which leads to the assembly of more even plant (i.e. weed in our case) communities composed of species that are poor competitors but highly stress tolerant (Goldberg & Novoplansky, 1997), which consequently have a reduced impact on crop productivity (Fig 5). On the contrary, in mild, productive environments, competition is stronger and more important; this leads to uneven weed communities, dominated by highly competitive weed species that are significantly detrimental to the crops.

## 5. Conclusions

Overall, our results demonstrate that intercropping alone does not reduce weed biomass or diversity, but that cereals play a crucial role in reducing weed pressure. In order to keep benefitting from the higher yield provided by diverse crop mixtures, we suggest that intercropped systems should include cereals. It is important to point out that for the highest level of crop diversity (i.e. 4-species mixtures), we could not assess the effect of crop diversity independently of the presence of each functional group. More research is therefore needed to investigate the effect of crop diversity on weed communities within these functional groups, as well as to understand which processes are responsible for the increased yield in more diverse crop mixtures. Our study also shows that crop– weed relationships are very dependent on local abiotic conditions, with the surprising result that crop yield did not correlate with weed suppression or weed diversity under harsher environmental conditions in Spain. This calls for further research on the assembly of weed communities and crop–weed relationships in resource-poor, low-productive environments, and further highlights the importance of developing locally adapted weed management strategies.

## Authors’ contributions

L.S. and C.S. conceived the study with input from N.E. L.S., N.E. and C.S. collected the data; L.S. assembled and analyzed the data with the help of C.S.; L.S. and C.S. wrote the paper. All authors discussed data analyses and results.

## Acknowledgments

We are grateful to Elisa Pizarro Carbonell, Carlos Barriga Cabanillas and Anja Schmutz for their help with the field experiment, and Johan Six for comments on the experimental design. We also thank the Aprisco de Las Corchuelas Field Station and the University of Zurich for the use of their facilities. The study was funded by the Swiss National Science Foundation (PP00P3_170645).

## Data accessibility

Data will be published on Zenodo upon publication of the manuscript.

## References

Adeux, G., Vieren, E., Carlesi, S., Bàrberi, P., Munier-Jolain, N., & Cordeau, S. (2019). Mitigating crop yield losses through weed diversity. Nature Sustainability, 2(11), 1018–1026.

Anderson, M. J. (2001). A new method for non-parametric multivariate analysis of variance. Austral Ecology, 26(1), 32–46.

Andrew, I. K. S., Storkey, J., & Sparkes, D. L. (2015). A review of the potential for competitive cereal cultivars as a tool in integrated weed management. Weed Research, 55(3), 239–248.

Bastiaans, L., Paolini, R., & Baumann, D. T. (2008). Focus on ecological weed management: What is hindering adoption? Weed Research, 48(6), 481–491.

Baumann, D. T., Kropff, M. J., & Bastiaans, L. (2000). Intercropping leeks to suppress weeds. Weed Research, 40(4), 359–374.

Bedoussac, L., Journet, E.-P., Hauggaard-Nielsen, H., Naudin, C., Corre-Hellou, G., Jensen, E. S., … Justes, E. (2015). Ecological principles underlying the increase of productivity achieved by cereal-grain legume intercrops in organic farming. A review. Agronomy for Sustainable Development, 35(3), 911–935.

Bertholdsson, N.-O. (2005). Early vigour and allelopathy - two useful traits for enhanced barley and wheat competitiveness against weeds. Weed Research, 45(2), 94–102.

Bertness, M. D., & Callaway, R. (1994). Positive interactions in communities. Trends in Ecology & Evolution, 9(5), 191–193.

Bilalis, D., Papastylianou, P., Konstantas, A., Patsiali, S., Karkanis, A., & Efthimiadou, A. (2010). Weed-suppressive effects of maize–legume intercropping in organic farming. International Journal of Pest Management, 56(2), 173–181.

Brainard, D. C., Bellinder, R. R., & Kumar, V. (2011). Grass–Legume Mixtures and Soil Fertility Affect Cover Crop Performance and Weed Seed Production. Weed Technology, 25(03), 473–479.

Brooker, R.W., Bennett, A.E., Cong, W.-F., Daniell, T.J., George, T.S., Hallett, P.D., … White, P.J. (2015), Improving intercropping: a synthesis of research in agronomy, plant physiology and ecology. New Phytol, 206, 107–117.

Brun, P., Zimmermann, N. E., Graham, C. H., Lavergne, S., Pellissier, L., Münkemüller, T., & Thuiller, W. (2019). The productivity-biodiversity relationship varies across diversity dimensions. Nature Communications, 10(1), 5691.

Bybee-Finley, K. A., Mirsky, S. B., & Ryan, M. R. (2017). Crop Biomass Not Species Richness Drives Weed Suppression in Warm-Season Annual Grass–Legume Intercrops in the Northeast. Weed Science, 65(5), 669–680.

Cierjacks, A., Pommeranz, M., Schulz, K., & Almeida-Cortez, J. (2016). Is crop yield related to weed species diversity and biomass in coconut and banana fields of northeastern Brazil? Agriculture, Ecosystems and Environment, 220, 175–183.

Connolly, J., Sebastià, M.-T., Kirwan, L., Finn, J. A., Llurba, R., Suter, M., … Lüscher, A. (2017). Weed suppression greatly increased by plant diversity in intensively managed grasslands: A continental-scale experiment. Journal of Applied Ecology, (August 2017), 852–862.

Corre-Hellou, G., Dibet, A., Hauggaard-Nielsen, H., Crozat, Y., Gooding, M., Ambus, P., … Jensen, E. S. (2011). The competitive ability of pea-barley intercrops against weeds and the interactions with crop productivity and soil N availability. Field Crops Research, 122(3), 264–272.

Duchene, O., Vian, J.-F., & Celette, F. (2017). Intercropping with legume for agroecological cropping systems: Complementarity and facilitation processes and the importance of soil microorganisms. A review. Agriculture, Ecosystems & Environment, 240, 148–161.

Goldberg, D., & Novoplansky, A. (1997). On the Relative Importance of Competition in Unproductive Environments. In Source: Journal of Ecology (Vol. 85).

Gomez, P., & Gurevitch, J. (1998). Weed community responses in a corn-soybean intercrop. Applied Vegetation Science, 1(2), 281–288.

Grime, J. P. (1973). Competitive exclusion in herbaceous vegetation. Nature, 242(5396), 344–347.

Grime, J. P. (1979). Plant strategies and vegetation processes. Plant Strategies and Vegetation Processes.

Hauggaard-Nielsen, H., Ambus, P., & Jensen, E. S. (2001). Interspecific competition, N use and interference with weeds in pea-barley intercropping. Field Crops Research, 70(2), 101–109.

Jabran, K., Mahajan, G., Sardana, V., & Chauhan, B. S. (2015, June 1). Allelopathy for weed control in agricultural systems. Crop Protection, Vol. 72, pp. 57–65.

Jensen, E. S., Carlsson, G., & Hauggaard-Nielsen, H. (2020). Intercropping of grain legumes and cereals improves the use of soil N resources and reduces the requirement for synthetic fertilizer N: A global-scale analysis. Agronomy for Sustainable Development, 40(1), 5.

Jesch, A., Barry, K. E., Ravenek, J. M., Bachmann, D., Strecker, T., Weigelt, A., … Scherer-Lorenzen, M. (2018). Below-ground resource partitioning alone cannot explain the biodiversity–ecosystem function relationship: A field test using multiple tracers. Journal of Ecology, 106(5), 2002–2018.

Lamb, E. G., & Cahill, J. F. (2008). When competition does not matter: grassland diversity and community composition. The American Naturalist, 171(6), 777–787.

Lamb, E. G., Kembel, S. W., & Cahill, J. F. (2009). Shoot, but not root, competition reduces community diversity in experimental mesocosms. Journal of Ecology, 97(1), 155–163.

Lange, M., Eisenhauer, N., Sierra, C. A., Bessler, H., Engels, C., Griffiths, R. I., … Scheu, S. (2015). Plant diversity increases soil microbial activity and soil carbon storage. Nat Commun, 6.

Legendre, P., & Andersson, M. J. (1999). Distance-based redundancy analysis: Testing multispecies responses in multifactorial ecological experiments. Ecological Monographs, 69(1), 1–24.

Liebman, M. (1988). Ecological suppression of weeds in intercropping systems: a review. Weed Management in Agroecosystems.

Mohler, C. L., & Liebman, M. (1987). Weed Productivity and Composition in Sole Crops and Intercrops of Barley and Field Pea. The Journal of Applied Ecology, 24(2), 685.

Mortensen, D., Bastiaans, L., & Sattin, M. (2000). The role of ecology in the development of weed management systems: an outlook. Weed Research.

Mouquet, N., Lagadeuc, Y., Devictor, V., Doyen, L., Duputié, A., Eveillard, D., … Loreau, M. (2015). Predictive ecology in a changing world. Journal of Applied Ecology, 52(5), 1293–1310.

Navas, M.-L. (2012). Trait-based approaches to unravelling the assembly of weed communities and their impact on agro-ecosystem functioning. Weed Research, 52(6), 479–488.

Oerke, E.-C., & Dehne, H.-W. (2004). Safeguarding production—losses in major crops and the role of crop protection. Crop Protection, 23(4), 275–285.

Ofori, F., & Stern, W. R. (1987). Cereal–Legume Intercropping Systems. Advances in Agronomy, 41(C), 41–90.

Pakeman, R. J., Brooker, R. W., Karley, A. J., Newton, A. C., Mitchell, C., Hewison, R. L., … Schöb, C. (2019). Increased crop diversity reduces the functional space available for weeds. Weed Research.

Palmer, M. W., & Maurer, T. A. (1997). Does diversity beget diversity? A case study of crops and weeds. Journal of Vegetation Science, 8(2), 235–240.

Petchey, O. L., & Gaston, K. J. (2002). Functional diversity (FD), species richness and community composition. Ecology Letters, (2002) 5: 402–411, 5, 402–411.

Poggio, S. L. (2005). Structure of weed communities occurring in monoculture and intercropping of field pea and barley. Agriculture, Ecosystems and Environment, 109(1–2), 48–58.

Randall, R. P. (2012). A Global Compendium of Weeds. Second Edition.

Schöb, C., Hortal, S., Karley, A. J., Morcillo, L., Newton, A. C., Pakeman, R. J., … Brooker, R. W. (2017). Species but not genotype diversity strongly impacts the establishment of rare colonisers. Functional Ecology, 31(7), 1462–1470.

Spehn, E. M., Hector, A., Joshi, J., Scherer-Lorenzen, M., Schmid, B., Bazeley-White, E., … Lawton, J. H. (2005). Ecosystem effects of biodiversity manipulations in european grasslands. Ecological Monographs, 75(1), 37–63.

Storkey, J., & Neve, P. (2018). What good is weed diversity? Weed Research, 58(4), 239–243.

Swanton, C. J., & Weise, S. F. (1991). Integrated Weed Management: The Rationale and Approach. Weed Technology, 5(3), 657–663.

Szumigalski, A., & Van Acker, R. (2005). Weed suppression and crop production in annual intercrops. Weed Science, 53(6), 813–825.

Tilman, D. (1982). Resource competition and community structure. Princeton University Press.

Tilman, D., Knops, J., Wedin, D., Reich, P., Ritchie, M., & Siemann, E. (1997). The influence of functional diversity and composition on ecosystem processes. Science, 277(5330), 1300–1302.

TimeTree :: The Timescale of Life, from http://timetree.org/

Tukey, J. W. (1957). On the Comparative Anatomy of Transformations. In Source: The Annals of Mathematical Statistics (Vol. 28).

Vandermeer, J. (1989). The Ecology of Intercropping.

Von Felten, S., Hector, A., Buchmann, N., Niklaus, P. A., Schmid, B., & Scherer-Lorenzen, M. (2009). Belowground nitrogen partitioning in experimental grassland plant communities of varying species richness. Ecology, 90(5), 1389–1399.

Worthington, M., & Reberg-Horton, C. (2013). Breeding Cereal Crops for Enhanced Weed Suppression: Optimizing Allelopathy and Competitive Ability. Journal of Chemical Ecology, 39(2), 213–231.

