## Supplementary Information for "Weeds are not always evil: crop-weed relationships are context-dependent and cannot fully explain the positive effects of intercropping on yield"

**Table S1.** List of crop species cultivars and their suppliers in Switzerland and in Spain.

| Species | Switzerland | | Spain | |
|---|---|---|---|---|
| | Cultivar | Supplier | Cultivar | Supplier |
| *Avena sativa* | Canyon | Sativa Rheinau | Previsión | INIA, Madrid |
| *Triticum aestivum* | Fiorina | DSP, Delley | Cabezorro (BGE015403) | INIA, Madrid |
| *Coriandrum sativum* | Indian | Zollinger Samen, Les Evouettes | wild type | Semillas Cantueso, Córdoba |
| *Chenopodium quinoa* | n.a. | Artha Samen, Münsingen | Atlas | Algosur, Sevilla |
| *Lupinus angustifolius* | Boregine | Aspenhof, Wilchingen | wild type | Semillas Cantueso, Córdoba |
| *Lens culinaris* | Anicia | Agroscope, Reckenholz | de la Armuña | Legumer SL, Salamanca |
| *Camelina sativa* | n.a. | Zollinger Samen, Les Evouettes | n.a. | Camelina Company, Madrid |
| *Linum usitatissimum* | Lirina | Sativa Rheinau | wild type | Semillas Cantueso, Córdoba |

**Table S2**. Results of mixed effects ANOVA testing environmental factors, ecotype, crop diversity and functional group presence on Weed Suppression Index.

*DenDF*, degrees of freedom of error term; *NumDF*, degrees of freedom of term; *F-value*, variance ratio; *Pr(>F)*, error probability. P-values in bold are significant at α = 0.05; * (P < 0.05), ** (P < 0.01), *** (P < 0.001).

| | *NumDF* | *DenDF* | *F value* | *Pr(>F)* | |
|---|---|---|---|---|---|
| *Country* | 1 | 16.19 | 241.1717 | **3.80E-11** | *** |
| *Fertilizer* | 1 | 17.63 | 47.8718 | **2.03E-06** | *** |
| *Crop species number* | 2 | 49.38 | 2.1044 | 0.132721 | |
| *Cereal* | 1 | 52.35 | 41.6284 | **3.60E-08** | *** |
| *Legume* | 1 | 48.23 | 0.2389 | 0.627235 | |
| *Superasterid herb* | 1 | 48.05 | 1.1541 | 0.28806 | |
| *Ecotype* | 1 | 471.64 | 11.2994 | **0.000838** | *** |
| *Country x fertilizer* | 1 | 15.7 | 7.367 | **0.015513** | * |
| *Country x crop species number* | 2 | 487.92 | 0.4047 | 0.667377 | |
| *Fertilizer x crop species number* | 2 | 462.82 | 0.4419 | 0.643115 | |
| *Country x ecotype* | 1 | 498.75 | 3.1275 | 0.077593 | |
| *Country x cereal* | 1 | 511.91 | 44.3065 | **7.24E-11** | *** |
| *Fertilizer x cereal* | 1 | 445.25 | 0.6053 | 0.436972 | |
| *Country x legume* | 1 | 510.89 | 11.3012 | **0.000833** | *** |
| *Fertilizer x legume* | 1 | 436.97 | 1.594 | 0.207436 | |
| *Country x superasterid herb* | 1 | 503.64 | 2.628 | 0.105621 | |
| *Fertilizer x superasterid herb* | 1 | 429.58 | 0.0982 | 0.75411 | |
| *Country x fertilizer x crop sp. number* | 2 | 479.62 | 1.1674 | 0.312061 | |
| *Country x fertilizer x cereal* | 1 | 500.19 | 10.3119 | **0.001407** | ** |

**Fig S1**. Effects of fertilization (a-b), ecotype (c-d), presence of a cereal (e-f), and legume (g) on weed species richness (left panel) and Shannon's diversity index (H) (right panel). (h): effect of country on Shannon's evenness index (J). (Abbreviations: CH = Swiss ecotype, ES = Spanish ecotype).

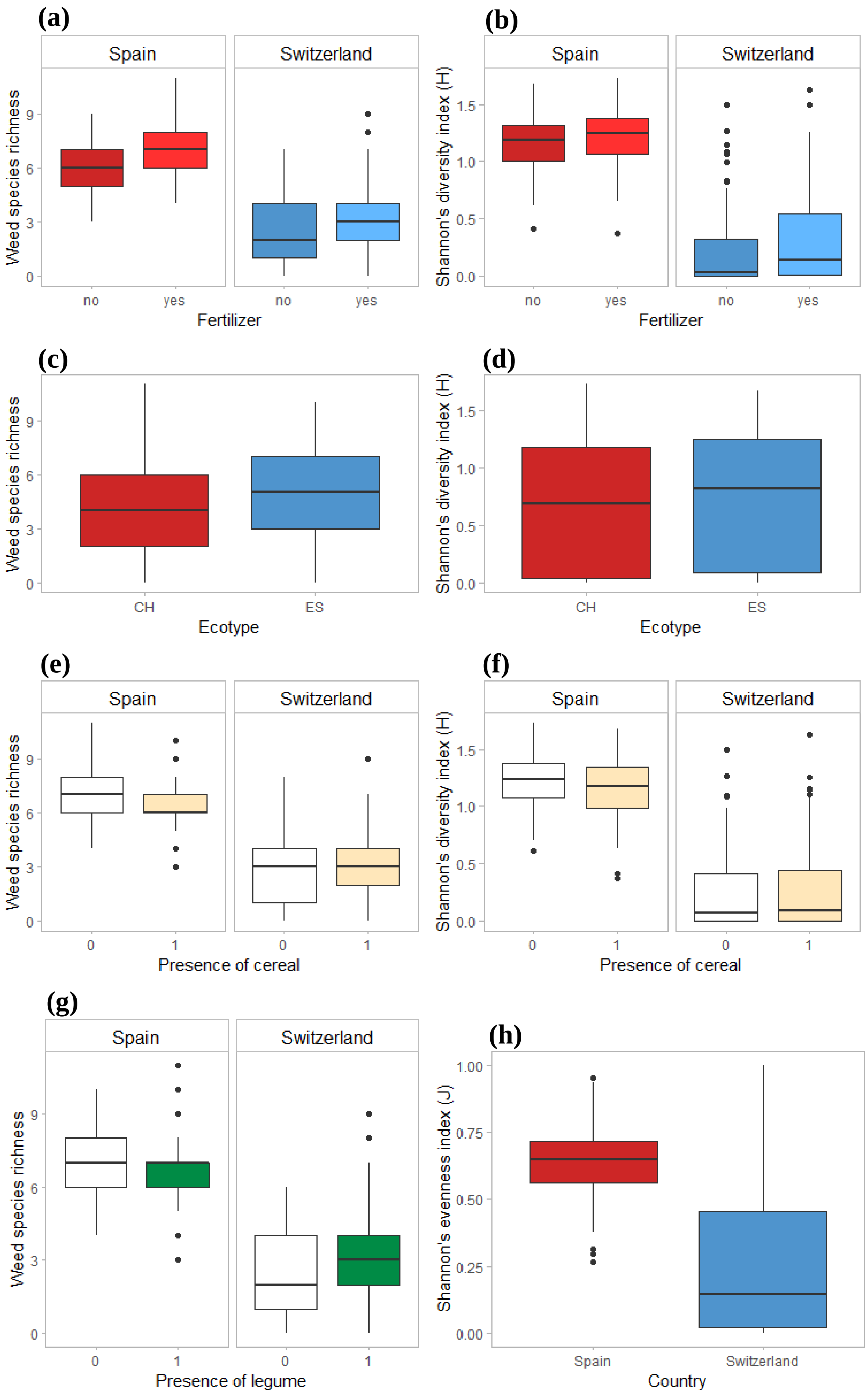

**Table S3**. Results of the permutational analyses of variance, showing $R^2$ and significance of the considered factors, in Switzerland (CH), and Spain.

P-values are significant at α = 0.05; * (P < 0.05), ** (P < 0.01), *** (P < 0.001).

| | *$R^2$ CH* | *Sign CH* | *$R^2$ Spain* | *Sign Spain* |
|---|---|---|---|---|
| *Fertilizer* | 0.01415 | *** | 0.15169 | *** |
| *Crop species number* | 0.00688 | | 0.01031 | * |
| *Cereal* | 0.00388 | | 0.14843 | *** |
| *Legume* | 0.00304 | | 0.00622 | * |
| *Superasterid herb* | 0.00334 | | 0.00572 | * |
| *Ecotype* | 0.00784 | ** | 0.02036 | *** |
| *Fertilizer x crop sp. number* | 0.00518 | | 0.00566 | |
| *Fertilizer x cereal* | 0.00481 | | 0.03218 | *** |
| *Fertilizer x legume* | 0.00157 | | 0.00179 | |
| *Fertilizer x superasterid herb* | 0.00516 | | 0.00297 | |
| *Crop sp. number x cereal* | 0.00224 | | 0.01038 | ** |
| *Crop sp. number x legume* | 0.00313 | | 0.00111 | |
| *Crop sp. number x superasterid herb* | 0.00260 | | 0.00287 | |

**Fig. S1** Constrained ordination plots showing changes in weed communities in response to crop species number × presence of cereal in Switzerland (a) and in Spain (b). The most responsive weed species are also represented (abbreviations: Po_an: *Poa annua;* St_me: *Stellaria media;* Pa_rh: *Papaver rhoeas;* Ar_th: *Arabidopsis thaliana;* Sp_ar: *Spergula arvensis;* Cap_bu: *Capsella bursa-pastoris;* Ec_cr: *Echinocloa crus-galli;* Ch_al: *Chenopodium album;* Am_re: *Amaranthus retroflexus;* Am_bl: *Amaranthus blithum;* So_ar: *Sonchus arvensis.*).

**(a)**

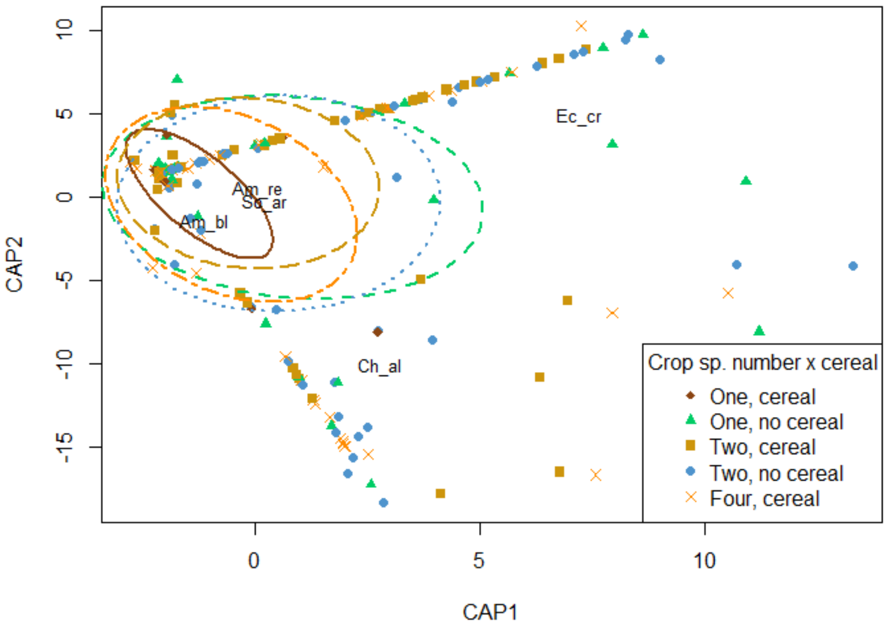


**(b)**

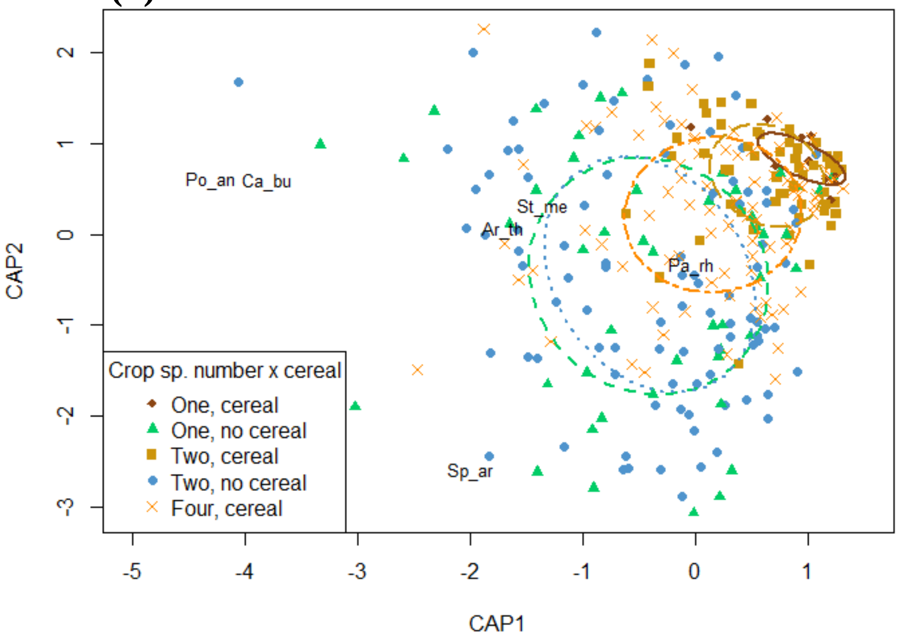

**Table S4**. Results of mixed effects ANOVA testing environmental factors, ecotype, crop diversity, functional group presence, Weed Suppression Index and weed species number on crop yield.

*DenDF*, degrees of freedom of error term; *NumDF*, degrees of freedom of term; *F-value*, variance ratio; *Pr(>F)*, error probability. P-values in bold are significant at α = 0.05; * (P < 0.05), ** (P < 0.01), *** (P < 0.001).

| | *NumDF* | *DenDF* | *F value* | *Pr(>F)* | |
|---|---|---|---|---|---|
| *WSI* | 1 | 117.6 | 29.6345 | **2.89E-07** | *** |
| *Weed species number* | 1 | 47.06 | 23.6461 | **1.33E-05** | *** |
| *Crop species number* | 2 | 40.54 | 4.6301 | **0.015451** | * |
| *Fertilizer* | 1 | 24.13 | 6.7191 | **0.015949** | * |
| *Country* | 1 | 67.21 | 0.1234 | 0.726491 | |
| *Ecotype* | 1 | 550.66 | 64.4422 | **6.04E-15** | *** |
| *Cereal* | 1 | 41.88 | 0.0148 | 0.903632 | |
| *Legume* | 1 | 40.1 | 1.6486 | 0.206525 | |
| *Superasterid herb* | 1 | 40.23 | 4.3005 | **0.044545** | * |
| *Country x WSI* | 1 | 406.59 | 24.1171 | **1.32E-06** | *** |
| *Fertilizer x WSI* | 1 | 202.19 | 0.3586 | 0.549951 | |
| *Country x weed sp. number* | 1 | 335.94 | 13.3182 | **0.000305** | *** |
| *Fertilizer x weed sp. number* | 1 | 79.99 | 4.0428 | **0.047729** | * |
| *Country x fertilizer* | 1 | 67.06 | 1.9833 | 0.163668 | |
| *Country x crop sp. number* | 2 | 533.1 | 11.8594 | **9.14E-06** | *** |
| *Fertilizer x crop sp. number* | 2 | 531.3 | 1.3957 | 0.248564 | |
| *Country x ecotype* | 1 | 543.14 | 3.4922 | 0.062196 | |
| *Country x cereal* | 1 | 545.7 | 17.6403 | **3.12E-05** | *** |
| *Fertilizer x cereal* | 1 | 538.12 | 0.023 | 0.879559 | |
| *Country x legume* | 1 | 546.54 | 147.6491 | **< 2.2e-16** | *** |
| *Fertilizer x legume* | 1 | 539.77 | 0.0131 | 0.909047 | |
| *Country x superasterid herb* | 1 | 544.62 | 0.9448 | 0.331472 | |
| *Fertilizer x superasterid herb* | 1 | 539.86 | 1.1486 | 0.284324 | |
| *Crop sp. number x WSI* | 2 | 559.91 | 1.5902 | 0.204799 | |
| *Country x fertilizer x crop sp. numb* | 1 | 555.57 | 0.0138 | 0.906442 | |
| *Country x fertilizer x WSI* | 2 | 529.46 | 0.1247 | 0.882747 | |
